# Wald’s martingale and the Moran process

**DOI:** 10.1101/2020.02.24.962407

**Authors:** Travis Monk, André van Schaik

## Abstract

Many models of evolution are stochastic processes, where some quantity of interest fluctuates randomly in time. One classic example is the Moran birth-death process, where that quantity is the number of mutants in a population. In such processes we are often interested in their absorption (i.e. fixation) probabilities, and the conditional distributions of absorption time. Those conditional time distributions can be very difficult to calculate, even for relatively simple processes like the Moran birth-death model. Instead of considering the time to absorption, we consider a closely-related quantity: the number of mutant population size changes before absorption. We use Wald’s martingale to obtain the conditional characteristic functions of that quantity in the Moran process. Our expressions are novel, analytical, and exact. The parameter dependence of the characteristic functions is explicit, so it is easy to explore their properties in parameter space. We also use them to approximate the conditional characteristic functions of absorption time. We state the conditions under which that approximation is particularly accurate. Martingales are an elegant framework to solve principal problems of evolutionary stochastic processes. They do not require us to evaluate recursion relations, so we can quickly and tractably obtain absorption probabilities and times of evolutionary stochastic processes.

**Author summary:** The Moran process is a probabilistic birth-death model of evolution. A mutant is introduced to an indigenous population, and we randomly choose organisms to live or die on subsequent time steps. Our goals are to calculate the probabilities that the mutant eventually dominates the population or goes extinct, and the distribution of time it requires to do so. The conditional distributions of time are difficult to obtain for the Moran process, so we consider a slightly different but related problem. We instead calculate the conditional distributions of the number of times that the mutant population size changes before it dominates the population or goes extinct. We use a martingale identified by Abraham Wald to obtain elegant and exact expressions for those distributions. We then use them to approximate conditional time distributions, and we show when that approximation is accurate. Our analysis outlines the basic concepts martingales and demonstrates why they are a formidable tool for studying probabilistic evolutionary models such as the Moran process.

## Introduction

The Moran birth-death process is a classic model of evolution [1, 2]. It considers a population of two species that we call mutants and indigenous. On sequential time steps, one organism is chosen to reproduce from either species. The newborn is the same species as its parent, and it replaces another organism in the population uniformly at random. The total population size then remains constant, but the numbers of mutant and indigenous organisms in it are stochastic in time. The mutants will drive the indigenous organisms to extinction with some probability after some random number of time steps. Determining these probabilities and times are principal problems in the analysis of the Moran process [3, 4] and its extensions [5–10].

The so-called ‘fixation probability’ of the Moran process is well-known [11–13]. However, the distribution of fixation time (and extinction time) is much harder to obtain. The means and variances of those distributions are known [5, 14, 15]. Their cumulants were recently reported, but only in the special cases of neutral selection or large population size [4]. They are known to obey a particular symmetry [16]. We have closed-form expressions for those conditional time distributions, but they are written as sums of products of eigenvalues of submatrices [3]. The definition of the Moran process is simple, but calculating its conditional time distributions is difficult.

Martingales can provide the full distribution of absorption times for a specific class of stochastic processes [17, 18]. Martingales are akin to conservation laws in physics. For example, in classical mechanics, we can invoke the conservation of energy to find the trajectory of a particle in a conserved system. Martingales are a statement that some quantity is conserved throughout a stochastic process. So if we know the value of that quantity at the beginning of the process, then we know it at the end. We can often calculate absorption (i.e. fixation) probabilities [8, 13] and/or times [17, 18] from it.

Abraham Wald identified a powerful martingale for stochastic processes whose steps are independent and identically distributed [17, 18]. Wald’s martingale is the seminal result of sequential analysis [19, 20]. From that martingale, he obtained absorption probabilities and the conditional characteristic functions (CFs) of absorption times. We will show that Wald’s martingale can be applied to the Moran process if we discard time steps where the mutant population size does not change. Therefore we can obtain the conditional CFs of the number of times that the mutant population size changes before it achieves fixation or goes extinct, i.e. ‘active steps’ [21]. We will then use those CFs to approximate the CFs of the time to fixation and extinction, and we will show the conditions under which that approximation is particularly accurate. Our CFs for the number of mutant population size changes are novel, clean, and exact results. More generally, Wald’s methodology demonstrates an elegant approach to investigate classic problems of evolutionary models [8, 13, 22].

## Results

Fig. 1 illustrates the problem considered by sequential analysis [19, 20].

**Fig 1.**
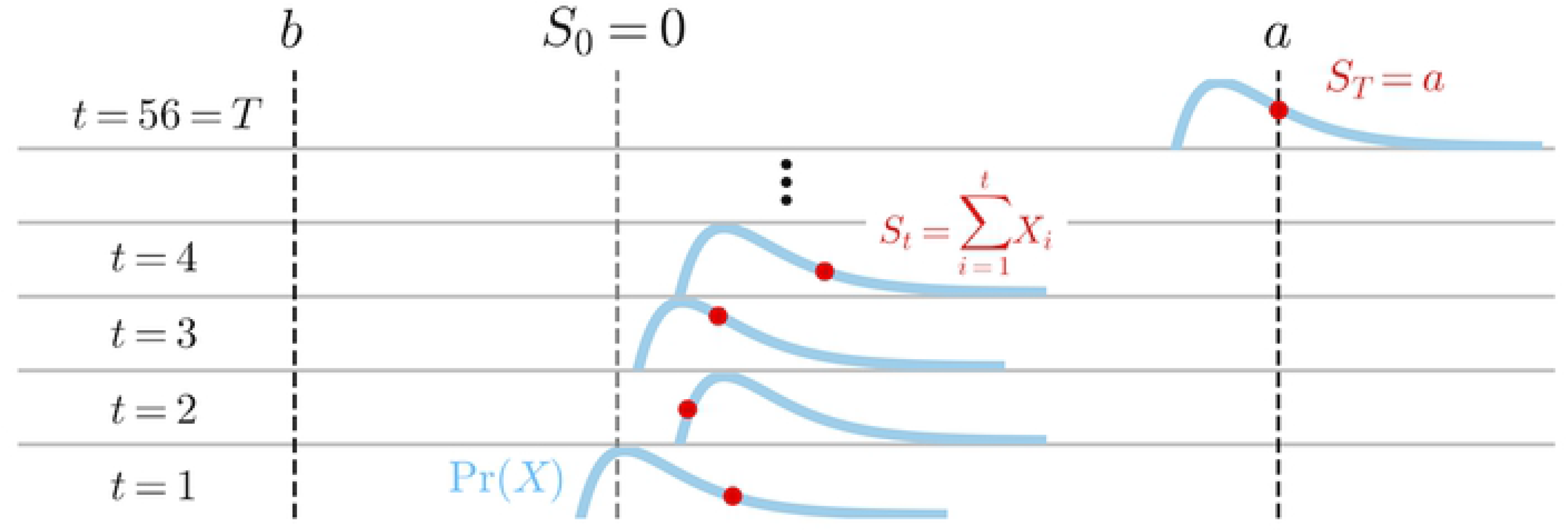
Schematic of sequential analysis. We initialize a cumulative sum to be *S*_0_ = 0. On subsequent time steps, we make one observation of *X* ~ Pr(*X*). The *X* are i.i.d. on each time step (blue curves). For illustrative purposes, we horizontally shift Pr(*X*) given *S*_*t*−1_ (red dots). In this example, the peak of the blue curve is horizontally aligned with the red dot below it. The sequential sum *S*_*t*_ of observations of *X* is a random walk in time *t*. While *S*_*t*_ remains between two absorbing barriers *a* and *b*, we continue adding observations of *X* to *S*_*t*_. We are interested in the probabilities that *S*_*t*_ hits either barrier first, and the distribution of the number of observations *T* it required to get there. In this example, *S*_*T*_ = *a*, and *T* = 56.

Let *S*_*t*_ be the cumulative sum of *t* realizations of independent and identically distributed (i.i.d.) random variables *X* ~ Pr(*X*) (Fig. 1 blue curves). Setting *S*_0_ = 0, we write 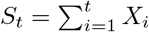. On sequential time steps, we observe one realization of *X* and add it to the sum of all previous realizations (Fig. 1, red dots). *S*_*t*_ is then a random walk in time.

Define two constant finite absorbing barriers *a* > 0 and *b* < 0 (vertical dashed lines, Fig. 1). While *b* < *S*_*t*_ < *a*, we continue making new observations of *X*, one per time step, and adding them to the sum. We stop this sequential process when *S*_*t*_ meets or exceeds either absorbing barrier (which happens in finite time [17]). We want to find the probability of hitting either barrier before the other, Pr(*S*_*T*_ = *a*) and Pr(*S*_*T*_ = *b*), i.e. ‘absorption probabilities.’ We also want to find the (conditional) distributions of the number of observations required to hit them, Pr(*T*|*S*_*T*_ = *a*) and Pr(*T*|*S*_*T*_ = *b*).

Wald derived these quantities from a martingale [17, 23, 24]:

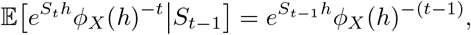

where *ϕ*_*X*_(*h*) is the moment generating function (MGF) of *X* and *h* is its independent variable. For the Moran process, *ϕ*_*X*_(*h*) exists because *X* can only assume the finite values +1, 0, or −1.

The quantity 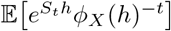 is conserved throughout a stochastic process whose steps are i.i.d. We illustrate this conservation by taking the expectation of both sides of Wald’s martingale:

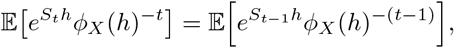

so by induction:

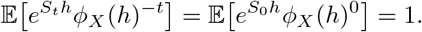

Doob’s optional stopping theorem [24, 25] states that a randomly-stopped martingale is still a martingale. Letting *T* be a random variable:

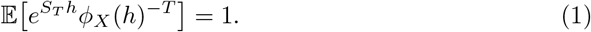

Eq. 1 is known as the fundamental identity in sequential analysis [17, 20]. It is a particularly powerful martingale because it is valid for all values of *h*. This property allows us to extract absorption probabilities and times from it by choosing special values of *h*.

Under weak assumptions, *ϕ*_*X*_(*h*) is convex, so it crosses 1 at two points *h* = 0 and *h* = *h*_0_ ≠ 0 (Lemma 2, [17]). We extract absorption probabilities by inserting *h* = *h*_0_ into Eq. 1:

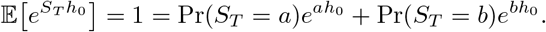

The process absorbs in finite time under weak conditions [17], so we can insert Pr(*S*_*T*_ = *b*) = 1 − Pr(*S*_*T*_ = *a*) and rearrange:

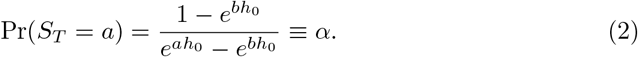

Wald [17] also showed that we can obtain the conditional CFs of *T* from his martingale. Splitting the expectation in Eq. 1:

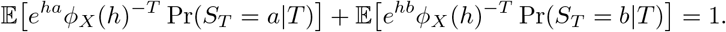

Since *ϕ*_*X*_(*h*) is convex (under weak assumptions [17]), its logarithm has two real roots in *h*, and its derivative is nonzero at those roots. Then for imaginary *τ*, − log *ϕ*_*X*_(*h*) = *τ* has two complex roots *h*_1_(*τ*) and *h*_2_(*τ*) in the neighborhood of *τ* = 0.

Inserting *ϕ*_*X*_(*h*_1_(*τ*)) = *e*^−*τ*^ and *ϕ*_*X*_(*h*_2_(*τ*)) = *e*^−*τ*^ in the expectations above, we obtain a system of two equations:

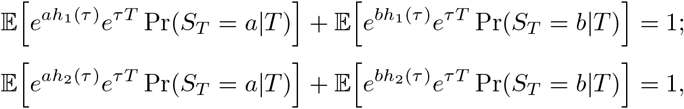

which can be written as conditional CFs of *T*, *ψ*_*T*|*b*_(*τ*) and *ψ*_*T*|*a*_(*τ*):

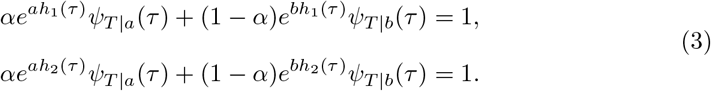

We have two equations, so we can solve for both conditional CFs.

Wald’s analysis is exact when *S*_*T*_ hits the absorbing barriers *a* and *b* exactly [17], as in the Moran process.

### Applying Wald’s martingale to the Moran process

For an introduction to the Moran process, see [12].

Consider the Moran birth-death process with a fixed population size *N*, and mutants with a relative reproductive advantage of *r* with respect to indigenous individuals [11–13]. Let *S*_*t*−1_ be the mutant population size on time step *t* − 1, and let *X*_*t*_ be the change in the mutant population size on time step *t*. Then 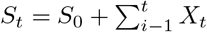, where *S*_0_ is the initial mutant population size.

Let *p*_*X*↑_, *p*_*X*↓_, and *p*_*X*0_ represent the Moran process transition probabilities, and let *F*_*t*−1_ = *rS*_*t*−1_ + *N* − *S*_*t*−1_ represent the total fitness of the graph on time step *t* − 1. The distribution of *X*_*t*_ is:

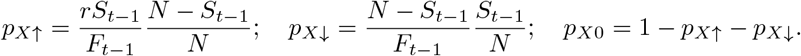

We cannot directly apply Wald’s martingale to the Moran process because the *X*_*t*_ are not independent. The distribution of *X*_*t*_ depends on the state *S*_*t*−1_, and therefore on previous observations of *X*.

Instead, consider the random variable *Y*_*t*_ ≡ *X*_*t*_|(*X*_*t*_ ≠ 0). In doing so we eliminate time steps where no change in the mutant population size occurs, i.e. we only consider ‘active steps’ [21]. Redefine 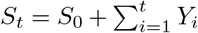, and let *p*_*Y*↑_ and *p*_*Y*↓_ represent transition probabilities:

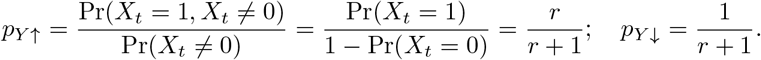

The transition probabilities of *Y*_*t*_ are independent and identical [4, 6]. Therefore Wald’s martingale is applicable to the Moran process if we eliminate time steps where the mutant population size does not change.

Eliminating time steps where *X*_*t*_ = 0 does not affect the fixation probability of the Moran process. We can show this by evaluating Eq. 2. First we find the value *h* = *h*_0_ ≠ 0 such that *ϕ*_*Y*_(*h*_0_) = 1:

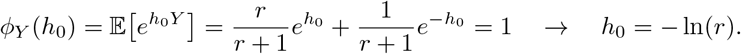

Typically we would set the Moran absorbing barriers to be *a* = *N* (fixation) and *b* = 0 (extinction), and we would set *S*_0_ to be some value between them. However, Wald’s analysis assumes that *S*_0_ = 0, so we shift the barriers to *a* = *N* − *S*_0_ and *b* = −*S*_0_. Inserting *h*_0_, *a*, and *b* into Eq. 2:

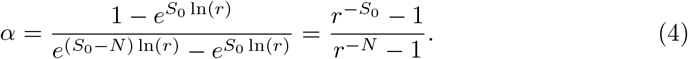

We recover the fixation probability of the Moran process [2, 11–13].

We can generalize Eq. 4 to find the probability that the Moran process achieves any state *a* before *b* starting from some *S*_0_ between them. Call that probability 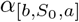:

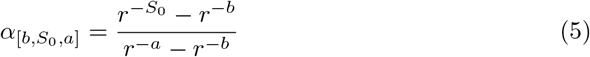

We next use Eq. 3 to find the conditional CFs of the number of mutant population size changes to achieve absorption *C*_*T*_. We begin by solving for the two roots *h*_1_(*τ*) and *h*_2_(*τ*) of *ϕ*_*Y*_(*h*) = *e*^−*τ*^:

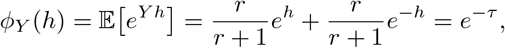

which has the analytical solution:

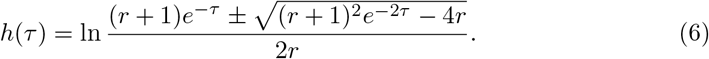

Fig. 2 plots these roots for the Moran process (Eq. 6) when *r* = .7 (top row) or *r* = 1.3 (bottom row). The left column plots the MGF of *Y*, which crosses 1 at *h* = 0 and *h*_0_ = − ln *r* (gray circles). The log of the MGF then has two complex roots *h*_1_(*τ*) (middle column) and *h*_2_(*τ*) (right column), where *τ* is purely imaginary. The real part (red) and imaginary part (black) of the roots are plotted separately. When *τ* = 0, the roots correspond to the two points where *ϕ*_*Y*_(*h*) = 1. Specifically, *h*_1_(0) = 0 + 0*i*, and *h*_2_(0) = *h*_0_ + 0*i*. These points are marked by the gray (imaginary part) and pink (real part) circles, respectively.

**Fig 2.**
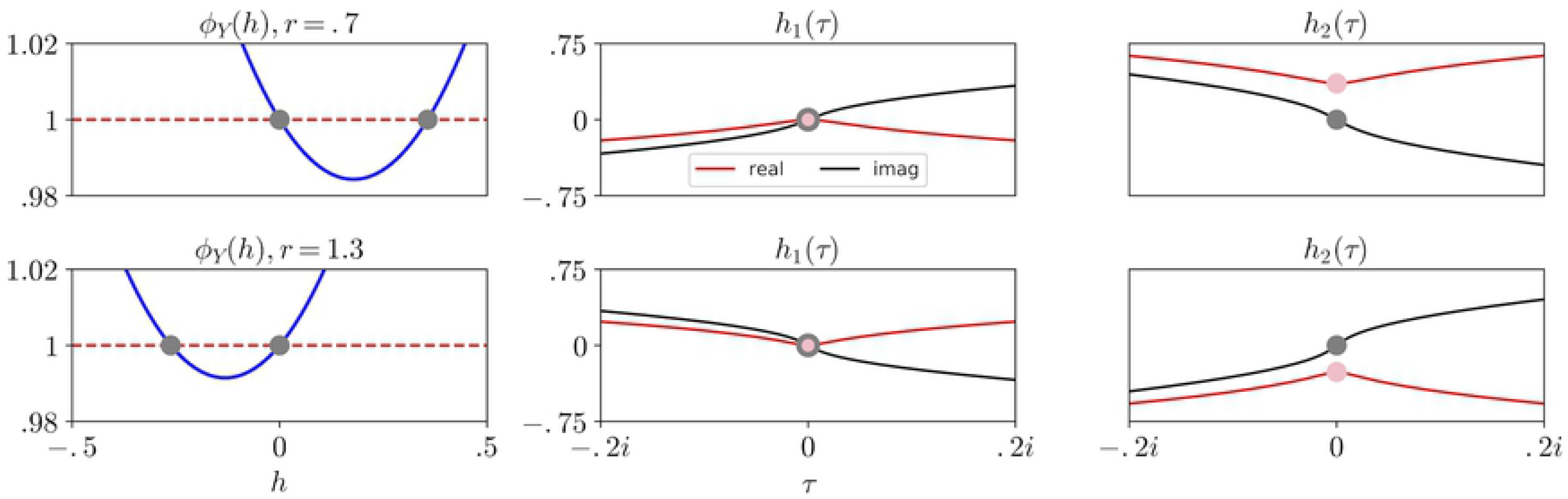
The roots *h*_1_(*τ*) and *h*_2_(*τ*) (Eq. 6) in the complex plane. Left column: *ϕ*_*Y*_(*h*) crosses 1 at two points for all *r* ≠ 1: *h* = 0 and *h* = − ln(*r*) (gray circles). Therefore its logarithm has two roots in the complex plane if *r* ≠ 1. Middle and right columns: real (red) and imaginary (black) parts of those roots *h*_1_(*τ*) and *h*_2_(*τ*), respectively. When *τ* = 0, the imaginary parts of all roots are 0 (gray circles). The real parts of *h*(0) are 0 (pink circles, middle column) or − ln(*r*) (pink circles, right column). Top row: *r* = 0.7. Bottom row: *r* = 1.3. Notice that the signs of the real and imaginary parts of both roots switch on either side of *r* = 1.

Fig. 2 shows that the signs of the real and imaginary parts of the roots depend on whether *r* < 1 or *r* > 1. We must take care in identifying which root is which. By convention, *h*_1_ is the root that passes through the origin of the complex plane. So if *r* < 1, then *h*_1_(*τ*) corresponds to subtraction and *h*_2_(*τ*) corresponds to addition of the ± sign in Eq. 6, and vice versa.

Fig. 3 shows the conditional CFs 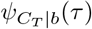 (left plots) and 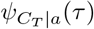 (right plots) that we obtain by inserting *h*_1_(*τ*) and *h*_2_(*τ*) into Eq. 3. We evaluated Eq. 3 for *r* = 0.9 (top plots) and *r* = 1.5 (bottom plots), and we used *N* = 10 and *S*_0_ = 3 in all plots. The real (pink) and imaginary (gray) parts of Eq. 3 are shown by the thick solid traces in Fig. 3. The real and imaginary parts of the CFs are even and odd, respectively, and they pass through 1 and 0 at *τ* = 0.

**Fig 3.**
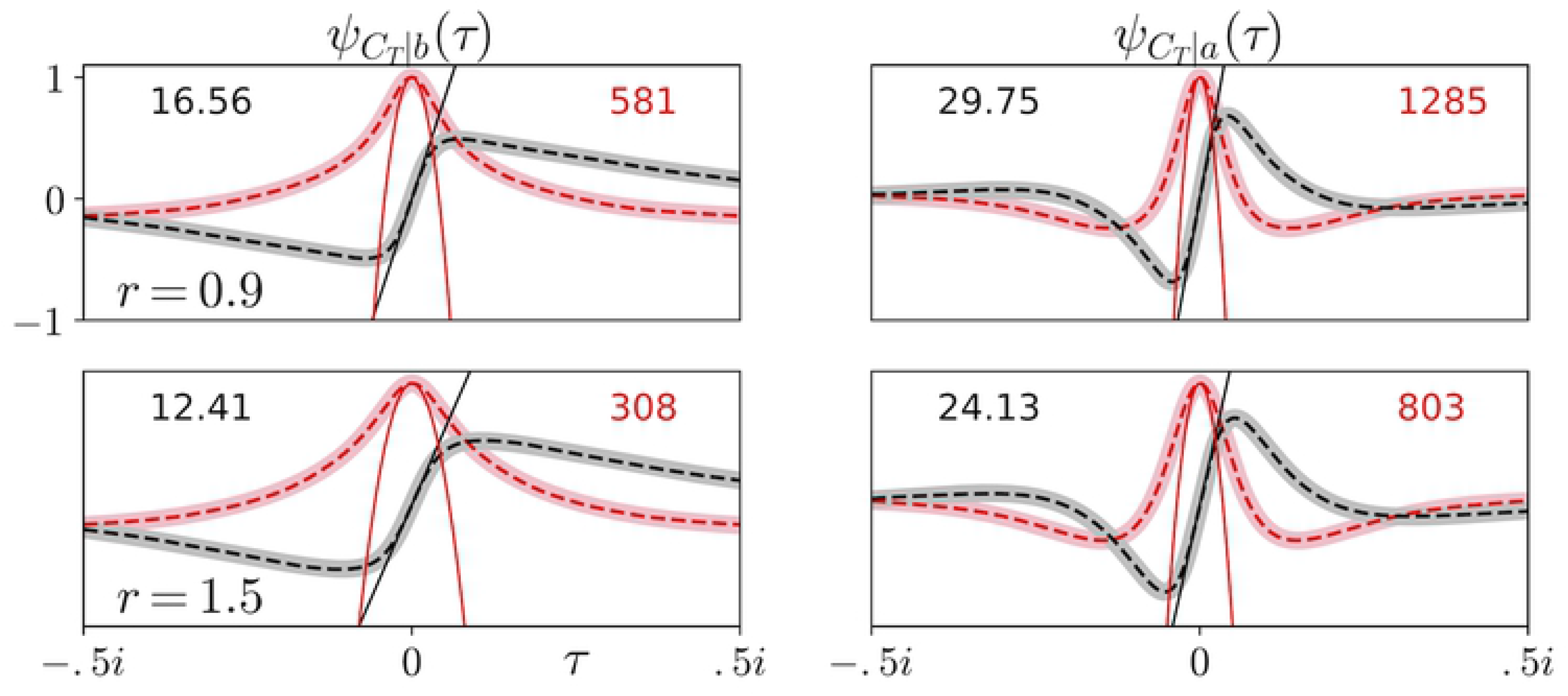
Conditional CFs for the number of changes in the mutant population size before absorption in the Moran process. CFs are conditional on the mutant population going extinct (left panels) or fixing (right panels). Real (pink) and imaginary (gray) parts are plotted separately. Fourier transforms of simulation results (red and black dashed lines) match theoretical predictions (thick pink and gray lines). The conditional first and second moments of *C*_*T*_ are visualized (black line and red parabola) and their values are reported (black and red numbers). Parameters were *S*_0_ = 3, *N* = 10, and *r* = .9 (top row) or *r* = 1.5 (bottom row).

Fig. 3 compares Eq. 3 with simulation results (dashed lines). We simulated the Moran process 100,000 times with the stated parameter values. We stored whether the initial mutant population fixed or went extinct, and how many times the mutant population size changed before doing so. We applied the Fourier transform to our simulated results to compare them with our predicted CFs. Their match is excellent because our analysis is exact, and we performed sufficiently many simulations for the Moran process to converge closely to that solution.

Notice that the conditional CFs in the top row of Fig. 3 are similar to those in the bottom row. Therefore the CFs of *C*_*T*_ are not particularly sensitive to changes in *r*. Comparing the left column with the right column, we see that the CFs of *C*_*T*_ are strongly influenced by where the Moran process absorbs. In particular, 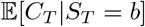 and 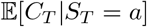 are given by the slopes of the imaginary parts of the CFs at *τ* = 0 (thin solid lines, Fig. 3). The black numbers in Fig. 3 report these slopes. For these parameter values, fixation requires almost double the number of mutant population size changes as extinction does on average. We also include a visualization of the second moment (red parabolas) and report its value (red numbers). The second moment increases as the mean increases, but notice that the second moment reduces when *r* is increased from 0.9 to 1.5 (c.f. red numbers, top and bottom row).

Fig. 4 plots conditional CFs 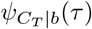 and 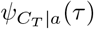 analogously to Fig. 3, except we vary *N* and *S*_0_ as shown instead of *r* (*r* = 1.3). We again verify that those conditional CFs (thick pink and gray lines) match Fourier transforms of 100,000 simulations as described above (thin dashed lines).

**Fig 4.**
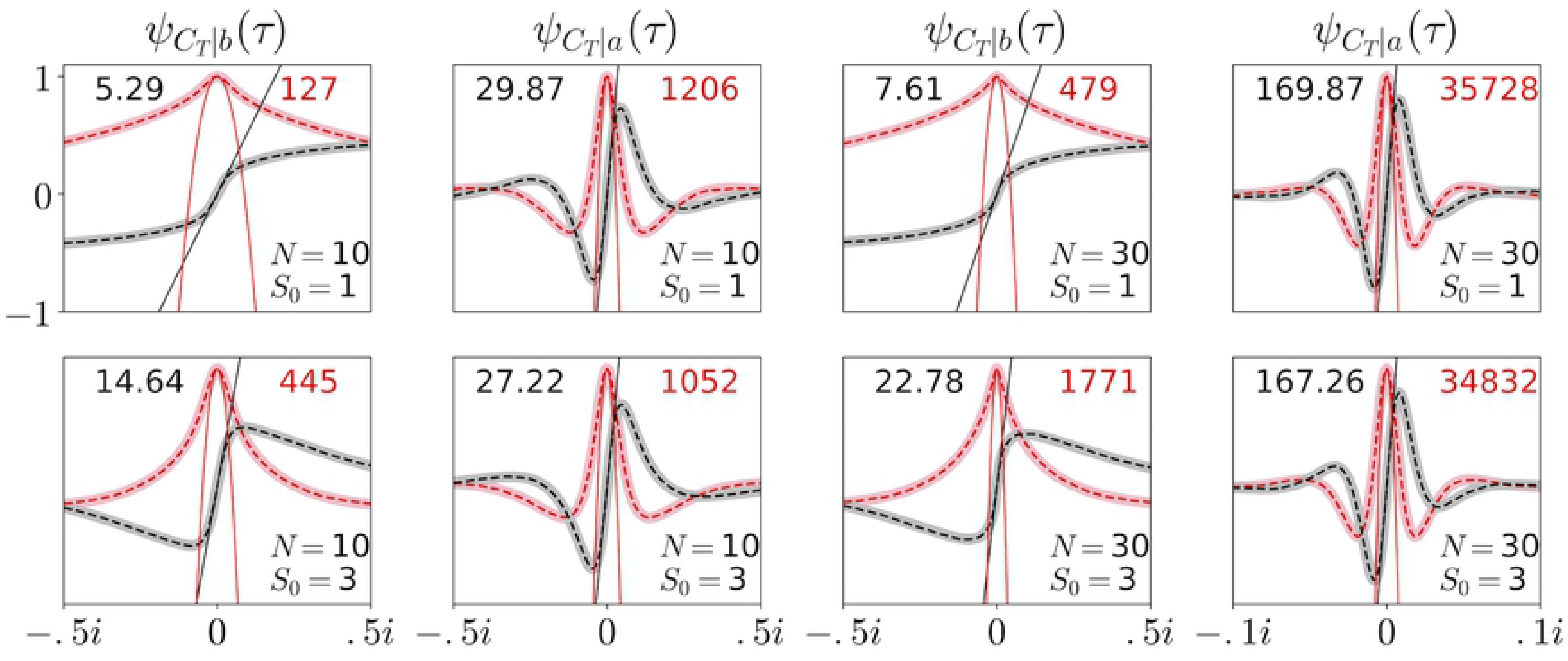
Conditional CF dependence on *N* and *S*_0_. Conditional CFs and simulation results plotted as in Fig. 3. *N* = 10 (left four plots) or 30 (right four plots), and *S*_0_ = 1 (top row) or 3 (bottom row). 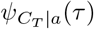 is sensitive to changes in *N* but not *S*_0_ if *S*_0_ ≪ *N* (the scale of the x-axis in the rightmost column is five times smaller than the others). 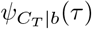 is sensitive to changes in *S*_0_ but not *N*. *r* = 1.3 in all plots. Simulations were run 100,000 times in each region of parameter space. The conditional first and second moments of *C*_*T*_ are visualized and their numbers are reported (thin solid black and red lines and numbers).

Fig. 4 shows that 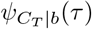 is not particularly sensitive to changes in *N*, but 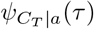 is (notice that the x-axis of the right column has a different scale than the other columns). We see that increasing *N* from 10 (left four plots) to 30 (right four plots) significantly increases the first two moments of *C*_*T*_|*S*_*T*_ = *a* (c.f. black and red numbers, second and fourth columns). This result reflects the fact that the Moran process requires more changes in the mutant population size to fix as the fixation barrier moves further from *S*_0_. Increasing *N* also increases 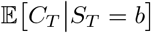 because longer paths to extinction become possible. For example, if *N* = 10, *S*_*t*_ cannot increase from *S*_0_ to 10 and then go extinct, because the mutant population already fixed. But if *N* = 30, then that path to extinction becomes possible (albeit unlikely if *r* > 1), and that path requires a large number of changes in the mutant population size.

Fig. 4 also shows that 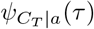 is not particularly sensitive to changes in *S*_0_, but 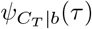 is. Increasing *S*_0_ from 1 (top row) to 3 (bottom row) significantly increases the first two moments of *C*_*T*_|*S*_*T*_ = *b* (c.f. black and red numbers, first and third column). Again, the Moran process requires more changes in the mutant population size to go extinct as the extinction barrier moves further from *S*_0_. Conversely, increasing *S*_0_ slightly reduces the first two moments of *C*_*T*_|*a*. However, slightly longer paths to fixation become possible if we increase *S*_0_ as well. For example, if *S*_0_ = 3, the mutant population can decrease to *S*_*t*_ = 1 and then achieve fixation. That path is slightly longer on average than a path that starts at *S*_0_ = 1 and achieves fixation. These two effects reduce each other’s impact on 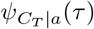 when *S*_0_ is small.

### Approximating the conditional CFs of absorption time

Fig. 5 shows that the conditional CFs of *C*_*T*_ can be used to accurately approximate the conditional CFs of *T*. Fig. 5 plots the conditional CFs of *C*_*T*_ with *r* = 1.3, *N* = 10, and *S*_0_ = 3 (the lower-left panel of Fig. 4 and its neighbor to the right). We also plot the Fourier transform of 100,000 simulations where we stored the conditional time *T* required to fix or go extinct (thin dashed lines). Real (pink and red) and imaginary (gray and black) parts are again plotted separately. We see that the conditional CFs of *C*_*T*_ and *T* are similar in form. The principal difference between them is the scale of *τ*. In Fig. 5, the bottom x-axes correspond to 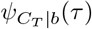 and 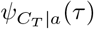, and the top x-axes correspond to *ψ*_*T*|*b*_(*τ*) and *ψ*_*T*|*a*_(*τ*).

**Fig 5.**
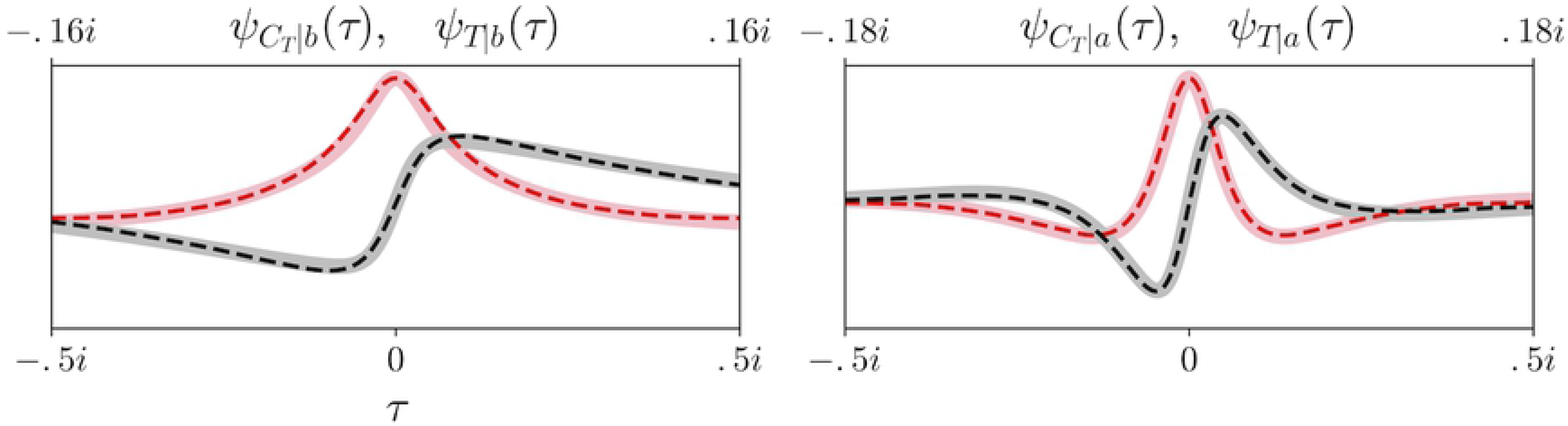
The conditional CFs of *C*_*T*_ can accurately approximate the conditional CFs of *T*. The main difference between them is the scale of *τ*. The conditional CFs of *C*_*T*_ are replotted from the two panels of Fig. 4 with *N* = 10 and *S*_0_ = 3. The thin dashed lines are Fourier transforms of 100,000 simulations of *T*|*b* (left panel) and *T*|*a* (right panel) using the same parameter values. Real (pink and red) and imaginary (gray and black) parts are plotted separately. Simulation results of *T* are plotted on a different scale of *τ* (top x-axis markings) than the theoretical results of *C*_*T*_ (bottom x-axis markings). If we can find those scaling factors *κ*_*a*_ (fixation) and *κ*_*b*_ (extinction), then we can approximate *ψ*_*T*|*a*_(*τ*) and *ψ*_*T*|*b*_(*τ*) from 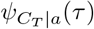 and 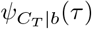.

This observation indicates that the conditional CFs of *T* can be approximated as:

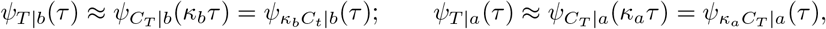

where *κ*_*b*_ and *κ*_*a*_ are scaling constants conditional on extinction or fixation, respectively.

A natural choice for *κ*_*b*_ and *κ*_*a*_ is the expected waiting time per change *T*_*C*_ in the mutant population size, conditional on fixation or extinction. Then we approximate *T* as (number of mutant population size changes) · (expected time per change):

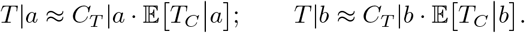

We now calculate 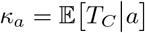 (the calculation of *κ*_*b*_ is analogous).

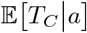 is the conditional expected time to absorption 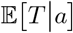 divided by the conditional expected number of mutant population changes 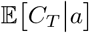.

The conditional expected time to fixation is:

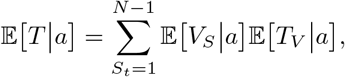

where *V*_*S*_ is the number of visits to state *S*_*t*_, and *T*_*V*_ is the time spent per visit [6].

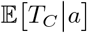 is then:

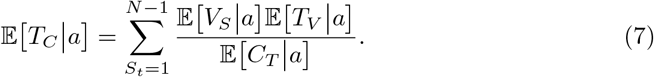

We now calculate the three conditional expectations in the summand.

#### The conditional expected number of visits to transient states

We calculate 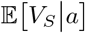 by noting that *V*_*S*_|*a* is geometrically distributed [26]. Let *p*_*V*_|*a* be the probability that the mutant population fixes without returning to the state *S*_*t*_ that it currently occupies. Let 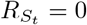 denote that occurrence. *p*_*V*_|*a* is then:

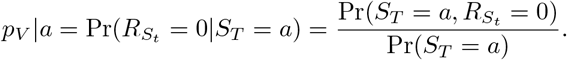

The only way that the Moran process can fix and not return to *S*_*t*_ is that a) the process increases by 1, and b) it then fixes without returning. We find these two absorption probabilities by inserting appropriate starting conditions and absorbing barriers into Eq. 5. For a) insert *S*_0_ = *S*_*t*_, *a* = *S*_*t*_ + 1, and *b* = *S*_*t*_ − 1, and for b) insert *S*_0_ = *S*_*t*_ + 1, *a* = *N*, and *b* = *S*_*t*_:

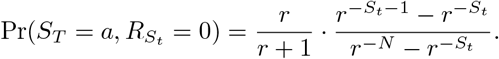

Pr(*S*_*T*_ = *a*) is the fixation probability (Eq. 4), but starting from *S*_0_ = *S*_*t*_. *p*_*V*_|*a* is then:

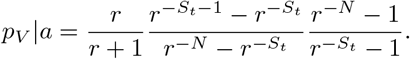

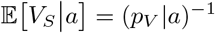, *if a state was visited at least once before fixation*. Since the Moran process fixes, we know that all states *S*_*t*_ ≥ *S*_0_ will be visited at least once. But for *S*_*t*_ < *S*_0_, we need the probability that the process arrives at state *S*_*t*_ at some time before it eventually fixes. Let *p*_*A*_|*a* represent that probability, and let 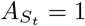 represent that occurrence:

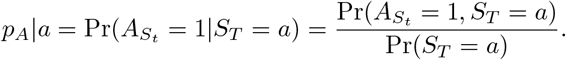

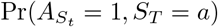 is the probability that the process, starting at *S*_0_, a) absorbs at *S*_*t*_ before *N*, and then b) absorbs at *N* before 0. For a) insert *S*_0_ = *S*_0_, *a* = *S*_*t*_, and *b* = *N*, and for b) insert *S*_0_ = *S*_*t*_, *a* = *N*, and *b* = 0 into Eq. 5. Pr(*S*_*T*_ = *a*) is given by Eq. 4. *p*_*A*_|*a* is then:

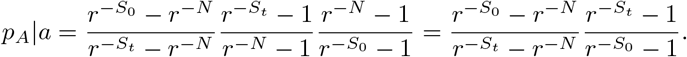

We can now find 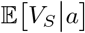:

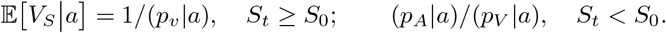

#### The conditional expected time spent in transient states per visit

We obtain 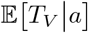 by noting that *T*_*V*_|*a* is also geometrically-distributed. Therefore 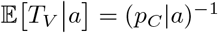, where *p*_*C*_|*a* is the probability that the mutant population size changes on a time step, conditional on fixation:

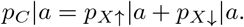

We find *p*_*X*↑_|*a* by Bayes’ rule, and we evaluate the likelihood by inserting *S*_0_ = *S*_*t*_ + 1, *a* = *N*, and *b* = 0 into Eq. 5:

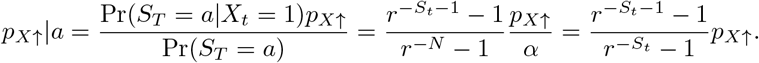

We find *p*_*X*↓_|*a* analogously:

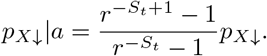

Inserting our expressions for *p*_*X*↑_ and *p*_*X*↓_:

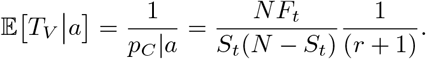

#### The conditional expected number of changes in the mutant population size

We can obtain 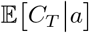 from 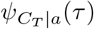, i.e. the slopes of the black lines in Figs. 3 and 4. We can also obtain it by summing 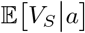 over all transient states:

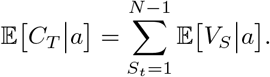

We can now insert 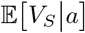, 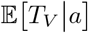, and 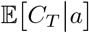 into Eq. 7 to obtain *κ*_*a*_. The calculations to obtain *κ*_*b*_ are analogous. We report them in the Jupyter notebook available as supporting information online.

Fig. 6 compares our approximation for *ψ*_*T*|*b*_(*τ*) and *ψ*_*T*|*a*_(*τ*) (thick solid lines) with Fourier transforms of simulation results (thin dashed lines). We ran 100,000 simulations of the Moran process for each of the displayed parameter settings and stored their absorption times conditional on fixation or extinction. Again, we plot the real and imaginary parts separately. Fig. 6 plots results from four regions of parameter space. In the left four plots, *r* = .95, and in the right four plots, *r* = 1.05. In the top row, *S*_0_ = 2, and in the bottom row, *S*_0_ = 8. *N* = 10 for all plots. We see that our theoretical approximations match simulation results reasonably closely in all plots.

**Fig 6.**
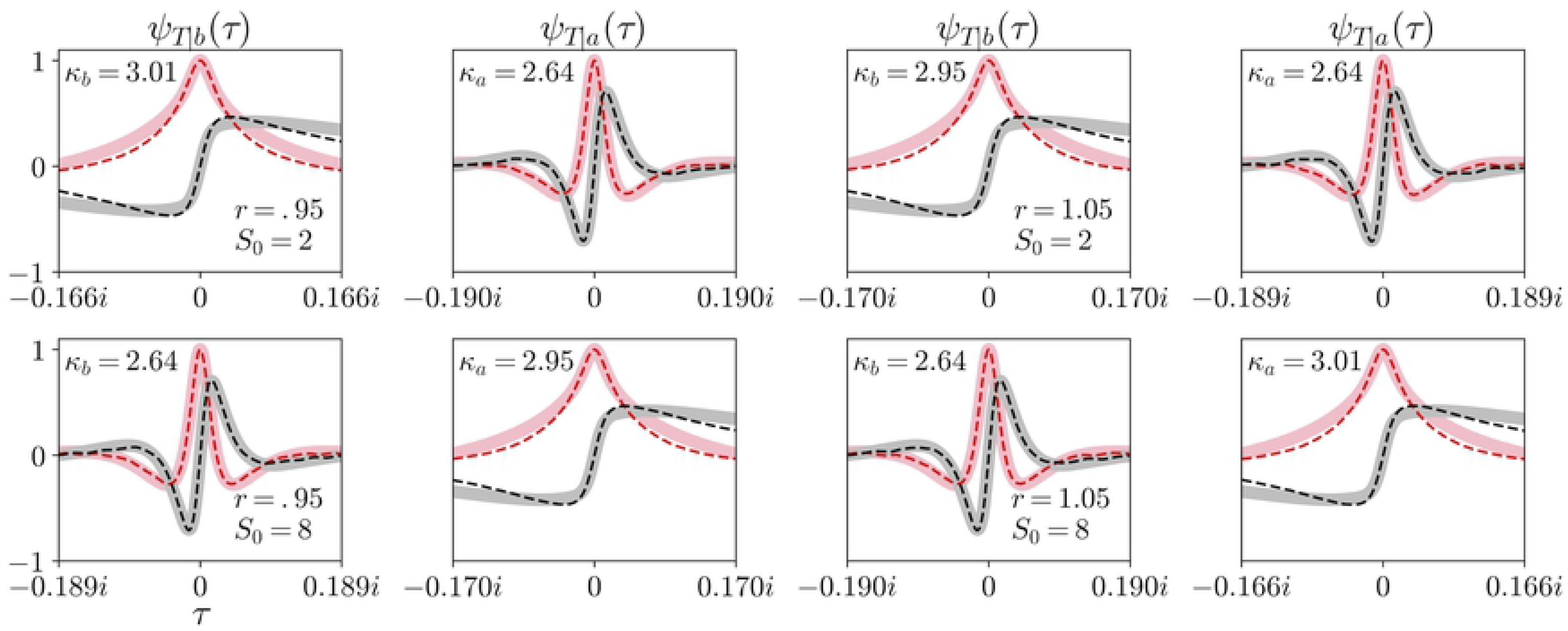
Investigating the accuracy of our approximation for the conditional CFs of *T*. Theoretical approximations (thick solid lines) and simulation results (thin dashed lines) are plotted analogously to Figs. 3 and 4. When *S*_0_ is low (top row), our approximation for *ψ*_*T*|*a*_(*τ*) is more accurate than it is for *ψ*_*T*|*b*_(*τ*). When *S*_0_ is high (bottom row), the converse is true. This observation holds whether *r* = .95 (left four plots) or *r* = 1.05 (right four plots). *N* = 10 for all plots. Scaling factors *κ*_*a*_ and *κ*_*b*_ are reported in each region of parameter space.

That match is closer in some plots than it is in others. The top row of Fig. 6 shows that, when *S*_0_ = 2, our approximation for *ψ*_*T*|*a*_(*τ*) matches simulation results very accurately. But our approximation for *ψ*_*T*|*b*_(*τ*) differs from simulation results noticeably, particularly further from the origin. The bottom row of Fig. 6 shows that, when *S*_0_ = 8, the converse is true.

### Sojourn CFs establish approximation accuracy

We can show why the accuracy of our approximation for conditional absorption times depends on *S*_0_ and where absorption occurs. Consider the conditional CFs 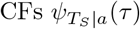 and 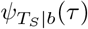 of the time *T*_*S*_ that the Moran process spends in each transient state; the ‘sojourn CFs’ [5, 16]. We can find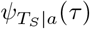 and 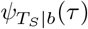 by noting that *T*_*S*_ = *V*_*S*_*T*_*V*_, i.e. the total time spent in a state is the number of visits to that state, multiplied by the time spent per visit. Since the *T*_*V*_ are i.i.d.:

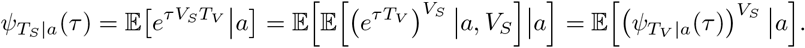

We can evaluate this conditional expectation. Using the tower property to split it, conditional on whether the process arrived at the transient state or not (i.e. *A* = 0 or *A* = 1):

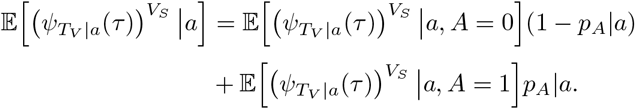

The first conditional expectation on the right hand side is 1, since *V*_*S*_ = 0 for states where the process never arrives. The second conditional expectation is a geometric sum that we can evaluate:

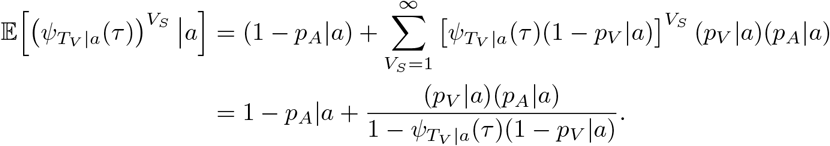

Finally, *T*_*V*_|*a* is also geometrically distributed, and the CF of a geometrically-distributed random variable is:

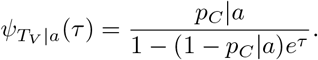

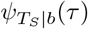 is calculated analogously.

Fig. 7 plots 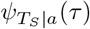 (first and third rows) and 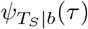 (second and fourth rows) for all transient states of the Moran process with *N* = 10 and *r* = .95. Again, we plot the real and imaginary parts separately. *S*_0_ = 2 (top two rows) or 8 (bottom two rows). The location of the black arrows in Fig. 7 indicate *S*_0_, and the direction that they point indicate whether the process fixed or went extinct. We compare our theoretical results with Fourier transforms of simulations (thin dashed traces), where we stored the amount of time that the Moran process spent in each transient state over 100,000 trials. Simulations and theory match extremely well because our sojourn CFs are exact, and we ran sufficiently many simulations of the Moran process to converge to them.

**Fig 7.**
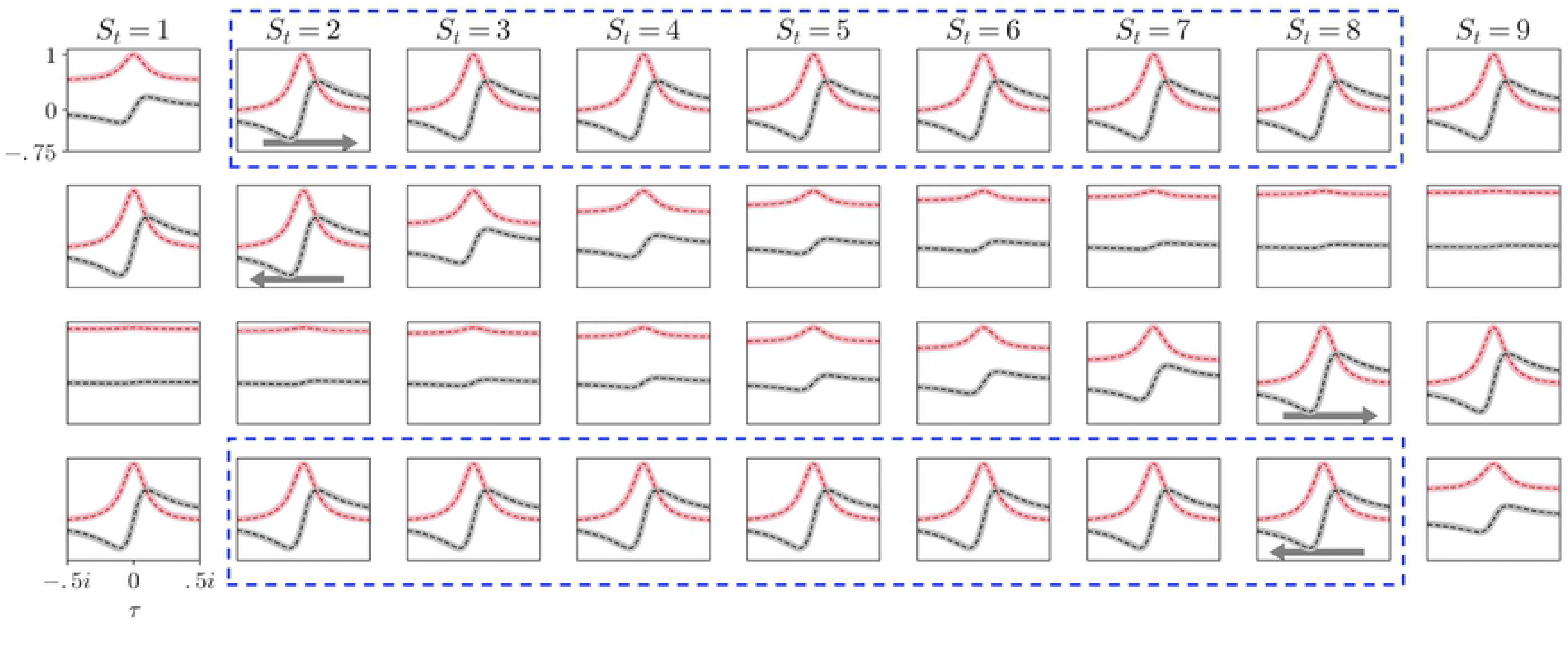
The sojourn CFs are relatively state-invariant when *S*_0_ is low and the process fixes, and vice versa. Conditional CFs of *T*_*S*_ (thick pink and gray lines) match perfectly with simulation results (thin dashed red and black lines). Location of arrows indicate *S*_0_ = 2 (top two rows) or 8 (bottom two rows). Direction of arrows indicate fixation (first and third rows) or extinction (second and fourth rows). *N* = 10 and *r* = .95 in all plots. When the conditional CFs are largely independent of *St*, we can accurately approximate *T*|*a* ∝ *C*_*T*|*a*_ and *T*|*b* ∝ *C*_*T*|*b*_.

Fig. 7 shows that when *S*_0_ = 2 and the Moran process fixes (top row), 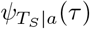 is almost identical for all transient states except *S*_*t*_ = 1. *T*|*a* is largely independent of which states the process occupies before fixation. Instead, *T*|*a* principally depends on how many state transitions occurred before fixation, i.e. *C*_*t*_|*a*. *T*|*a* and *C*_*t*_|*a* are then approximately proportional to each other, so our approximation is accurate. The same observation is true when *S*_0_ = 8 and the Moran process goes extinct (Fig. 7 bottom row). When *S*_0_ = 2 and the Moran process goes extinct (Fig. 7 second row), 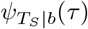 differs substantially among *S*_*t*_. *T*|*b* then depends not only on *C*_*T*_|*b* but also on which states the process occupied, and how many times each state was visited, before extinction. Our approximation *T*|*b* ∝ *C*_*T*_|*b* will be less accurate because *T*|*b* is sensitive to the specific path to extinction, as well as the length of the path (i.e. the value of *C*_*T*|*b*_). Collectively, these results suggest that we can accurately approximate *ψ*_*T*|*a*_(*τ*) when *S*_0_ is small, or *ψ*_*T*|*b*_(*τ*) when *S*_0_ is large.

Notice that some sojourn CFs in Fig. 7 appear very similar to each other. In fact, the sojourn CFs in the dashed blue boxes of Fig. 7 vary slightly between transient states, but 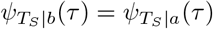 for each state 2 ≤ *S*_*t*_ ≤ 8. This identity arises because *p*_*V*_|*b* = *p*_*V*_|*a*, and *p*_*C*_|*b* = *p*_*C*_|*a* for all transient states (see Jupyter notebook, supporting material online). The sojourn CFs 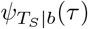 and 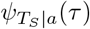 only differ from each other via the probabilities of arriving at a state, *p*_*A*_|*a* and *p*_*A*_|*b*. So in the special case when *S*_0_ = 1 and the Moran process fixes, or when *S*_0_ = *N* − 1 and the process goes extinct, the process will arrive at every state. The only asymmetry distinguishing 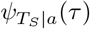 and 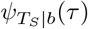 is then eliminated, and the conditional sojourn CFs equal each other for all transient states. Therefore the conditional time distributions of fixation from *S*_0_ = 1 and extinction from *S*_0_ = *N* − 1 are equivalent, as reported in [16].

All figures and simulations can be reproduced with a Jupyter notebook available as supporting material online.

## Discussion

The Moran process was first introduced over 60 years ago, yet its exact conditional distributions of the time to absorption remain an open research topic [3, 4]. Those distributions are difficult to evaluate because the amounts of time that the Moran process spends in each transient state are interdependent. For example, say the state *S*_*t*_ = 5 was visited *V*_5_ = 100 times. Then we know that states *S*_*t*_ = 4 and *S*_*t*_ = 6 are less likely to have been visited once or twice than if we did not have that information. Since the number of visits to a state depend on the number of visits to neighboring states, so do the amounts of time that the process spends in them.

When faced with such a challenging problem as calculating conditional time distributions, we have a few options. First, we can either make simplifying approximations to the problem [4]. Second, we can settle for closed-form but intractable expressions for those distributions [3] and numerically evaluate them. Third, we can tweak the problem itself. Rather than calculate *T*, we can exactly and easily calculate *C*_*T*_, and observe that those distributions are related to each other. The conditional, general, and exact distribution of *T* might be difficult to obtain, but we can get a tractable proxy for it.

When they are applicable, martingales have several advantages compared to other popular approaches to analyzing evolutionary stochastic processes. Markov chain analysis seems to be the predominant method for studying these kinds of problems in the literature [4, 27–36]. The Markov chain approach constructs a matrix of transition probabilities and calculates e.g. absorption probabilities and times from it. While Markov chains are general and flexible tools, those calculations require us to evaluate recursions over all state space. Rarely, those recursions can be evaluated inductively [27, 31]. We can calculate them by brute force if the size of state space is small enough [5, 31, 32, 37]. But quite often they are written using sums of products over all state space [4, 5, 33–36]. If we consider more complicated evolutionary stochastic processes, e.g. an evolutionary graph with many partitions [8, 11, 38], Markov chains quickly become an infeasible method of analysis. Martingales circumvent these issues because they are conservation statements, so we do not need to evaluate recursions over state space.

Other popular approaches to studying evolutionary stochastic processes include simulation [34, 39–41] and approximation [11, 37, 42, 43]. While both approaches are useful, they also have drawbacks. In principle, any well-defined stochastic process can be studied by simulating it enough. But simulation results cannot be generalized beyond its specific parameter settings. Computation time can prohibit us from evaluating a stochastic process over all parameter space, particularly if the dimensionality of parameter space is high. Approximations can yield tractable analytical expressions for quantities of interest, but they require us to take a special limit such as large population size [11] or weak selection [37, 42, 43]. Such results are only valid in those particular limits. Martingales can provide exact results that are valid over all parameter space without requiring simplifying assumptions.

The key to employing the martingale methodology is to find a quantity that is independent of the state of an evolutionary stochastic process. That quantity is then conserved throughout the process, so we can use Doob’s optional stopping theorem [24, 25] to extract statistics of interest. There are a couple of ways to find such quantities. We can modify the process such that transition probabilities are state-independent, as we have done here by discarding time steps where the mutant population size remains unchanged. Alternatively, we can show that the expectation of some state-dependent quantity always equals 1 (a product martingale) or 0 (a sum martingale), regardless of the state [8, 13]. Finding such a quantity can be difficult, in which case we should revert to other methodologies such as Markov chain analysis, simulation, or approximation. But if we can find such a quantity, then we can quickly obtain elegant expressions for the global statistics of an evolutionary process. So other methodologies such as Markov chains, simulations, and approximations should not be our default approaches to studying such problems, but rather our fallback options.

